# Comparative untargeted metabolomic profiling of induced mitochondrial fusion in pancreatic cancer

**DOI:** 10.1101/2021.08.06.455450

**Authors:** Nicholas D. Nguyen, Meifang Yu, Vinit Y. Reddy, Ariana Acevedo-Diaz, Enzo C. Mesarick, Joseph Abi Jaoude, Min Yuan, John M. Asara, Cullen M. Taniguchi

**Author notes:** These authors contributed equally to this work (Nicholas D. Nguyen, Meifang Yu). To whom correspondence should be addressed: Cullen M. Taniguchi, MD, PhD, The University of Texas MD Anderson Cancer Center, Division of Radiation Oncology, 1515 Holcombe Blvd., Unit 1050, Houston, TX 77030-4000.

## Abstract

Mitochondria are dynamic organelles that constantly alter their shape through the recruitment of specialized proteins, like mitofusin-2 (Mfn2) and dynamin-related protein 1 (Drp1). Mfn2 induces the fusion of nearby mitochondria, while Drp1 mediates mitochondrial fission. We previously found that the genetic or pharmacological activation of mitochondrial fusion was tumor suppressive against pancreatic ductal adenocarcinoma (PDAC) in several model systems [1]. The mechanisms of how these different inducers of mitochondrial fusion reduce pancreatic cancer growth are still unknown. Here, we characterized and compared the metabolic reprogramming of these three independent methods of inducing mitochondrial fusion in KPC cells: overexpression of Mfn2, genetic editing of Drp1, or treatment with leflunomide. We identified significantly altered metabolites via robust, orthogonal statistical analyses and found that mitochondrial fusion consistently produces alterations in the metabolism of amino acids. Our unbiased methodology revealed that metabolic perturbations were similar across all these methods of inducing mitochondrial fusion, proposing a common pathway for metabolic targeting with other drugs.

## Introduction

Pancreatic ductal adenocarcinoma (PDAC) relies on mitochondrial respiration through remodeling of the electron transport chain in order to sustain its proliferative abilities [2,3]. We and others have found that morphological changes in mitochondria can alter their function [1,4]. Mitochondria undergo fusion and fission in response to external stimuli to optimize metabolic functions and to promote turnover of damaged organelles through mitophagy [5]. This balance between mitochondrial fusion and fission is regulated by two key molecules: mitofusin-2 (Mfn2) and dynamin-related protein 1 (Drp1). As its name suggests, Mfn2 directs the fusion of outer membranes in adjacent mitochondria whereas Drp1 aggregates to the surface of elongated networks, constricting the mitochondrial membranes until they break apart through the process of mitochondrial fission.

Pancreatic cancer cells often display aberrations of mitochondrial dynamics in favor of mitochondrial fission [1,6], where these organelles take on a fragmented appearance, which appears to be a KRAS-dependent phenomenon [7]. We previously demonstrated that this overactive mitochondrial fission could be therapeutically targeted by disrupting Drp1, increasing expression of Mfn2 genetically, or with the use of leflunomide [1]. The net effect of these three interventions promoted mitochondrial fusion, which curbed oxidative phosphorylation (OXPHOS), thereby suppressing tumor growth in pancreatic cancer [1]. However, the metabolic mechanism by which mitochondrial fusion reduced PDAC growth is still unknown.

To understand how mitochondrial fusion alters the cellular metabolism of PDAC, we performed an unbiased comparative metabolomic analysis between three different methods of inducing mitochondrial fusion: (1) genetically inducing fusion using a tetracycline-inducible system to overexpress Mfn2 (Tet-On Mfn2), (2) directly inhibiting the decomposition of mitochondrial fusion through genetically knocking out Drp1 using CRISPR (sgDrp1), and (3) pharmacologically inducing fusion through treatment with leflunomide. We found common metabolic pathways between these different methods of inducing mitochondrial fusion, suggesting areas for metabolic intervention to further optimize this therapeutic target.

## Results

### Mitochondrial Fusion Distinctly Alters PDAC Metabolome

We created isogenic cell lines derived from murine KPC pancreatic tumors with proven mitochondrial fusion [8]. Tet-On Mfn2 cells express Mfn2 upon exposure to low-doses of doxycycline, which induces mitochondrial fusion compared to doxy-negative controls. We also produced cells with predominantly fused mitochondria by CRISPR-mediated abrogation of Drp1 or treatment with Leflunomide (Figure 1). We extracted metabolites from each cell line in a minimum of five biological replicates along with isogenic controls and subjected them to mass spectrometric analysis after steady state metabolite collection using a well-established methanol extraction method [9,10].

**Figure 1.**
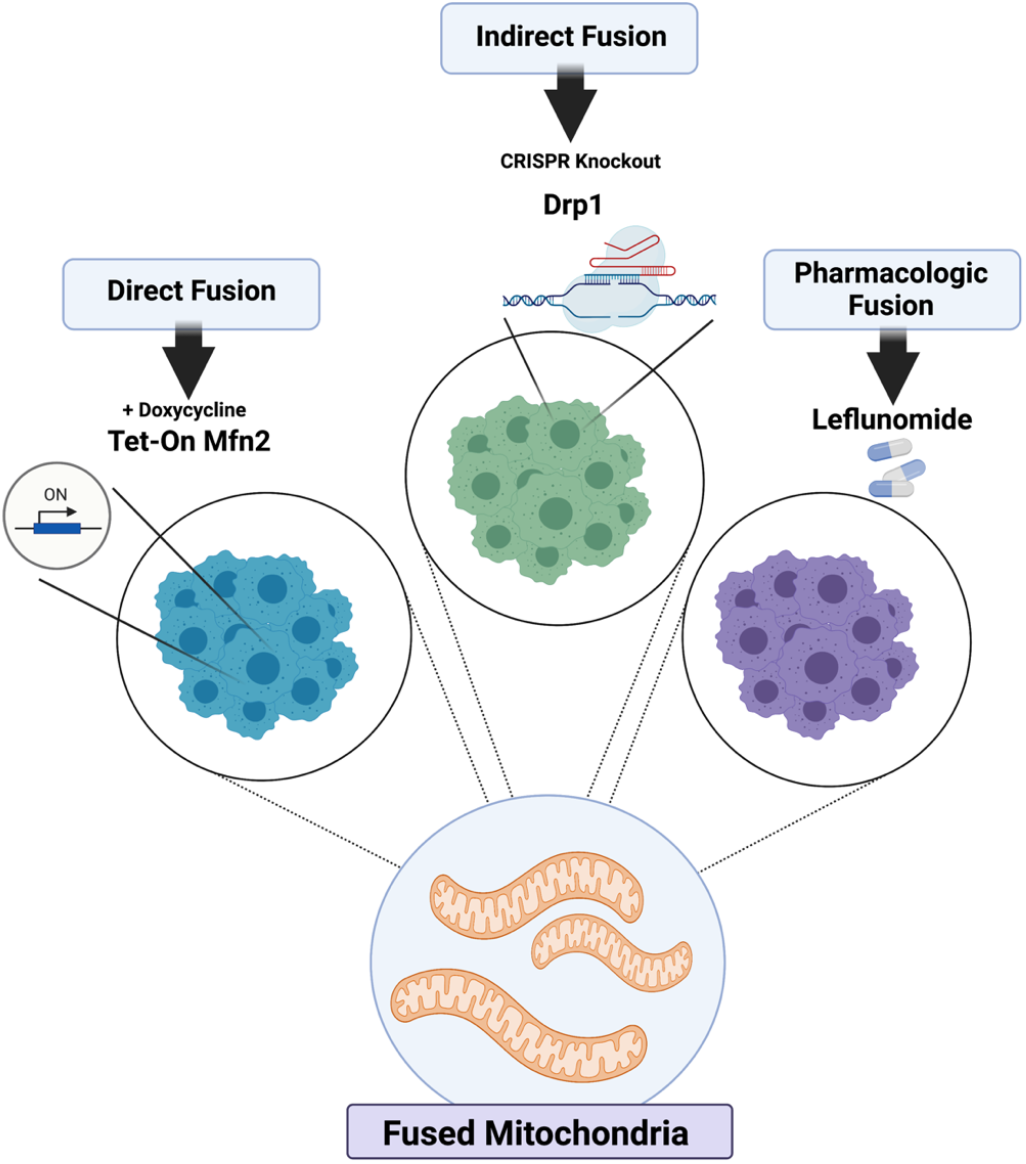
Models of mitochondrial fusion induction. KPC cells were genetically modified to directly overexpress Mfn2 in a tetracycline-inducible manner, indirectly fuse through CIRSPR knockout of Drp1, and pharmacologically fuse after treatment with Leflunomide.

Liquid chromatography tandem mass spectrometry (LC-MS/MS) analysis quantified the relative concentration levels of 296 distinct metabolites for each of the cells. To understand the metabolites more closely correlated with mitochondrial fusion, we subjected the data to stringent filters and normalized the datasets to the control of each experimental group. Supervised partial least-squares discriminant analysis (PLS-DA) and unsupervised principal component analysis (PCA) revealed distinct clustering between the induced mitochondrial fusion groups (n = 6) and their respective controls (n = 6, Figure 2A and 2B). Further hierarchical clustering using a euclidean distance and ward clustering algorithm revealed that individual replicates for each treatment group clustered together (Figure 2C). Overall, a total of 14 significantly altered metabolism pathways were shared between the Tet-On Mfn2, sgDrp1, and pharmacologic treated Leflunomide groups (Table 1). Metabolic super-pathways regulating amino acids and nucleotides were most consistently altered by the induction of mitochondrial fusion, compared to individual controls (Figure 2C).

**Figure 2.**
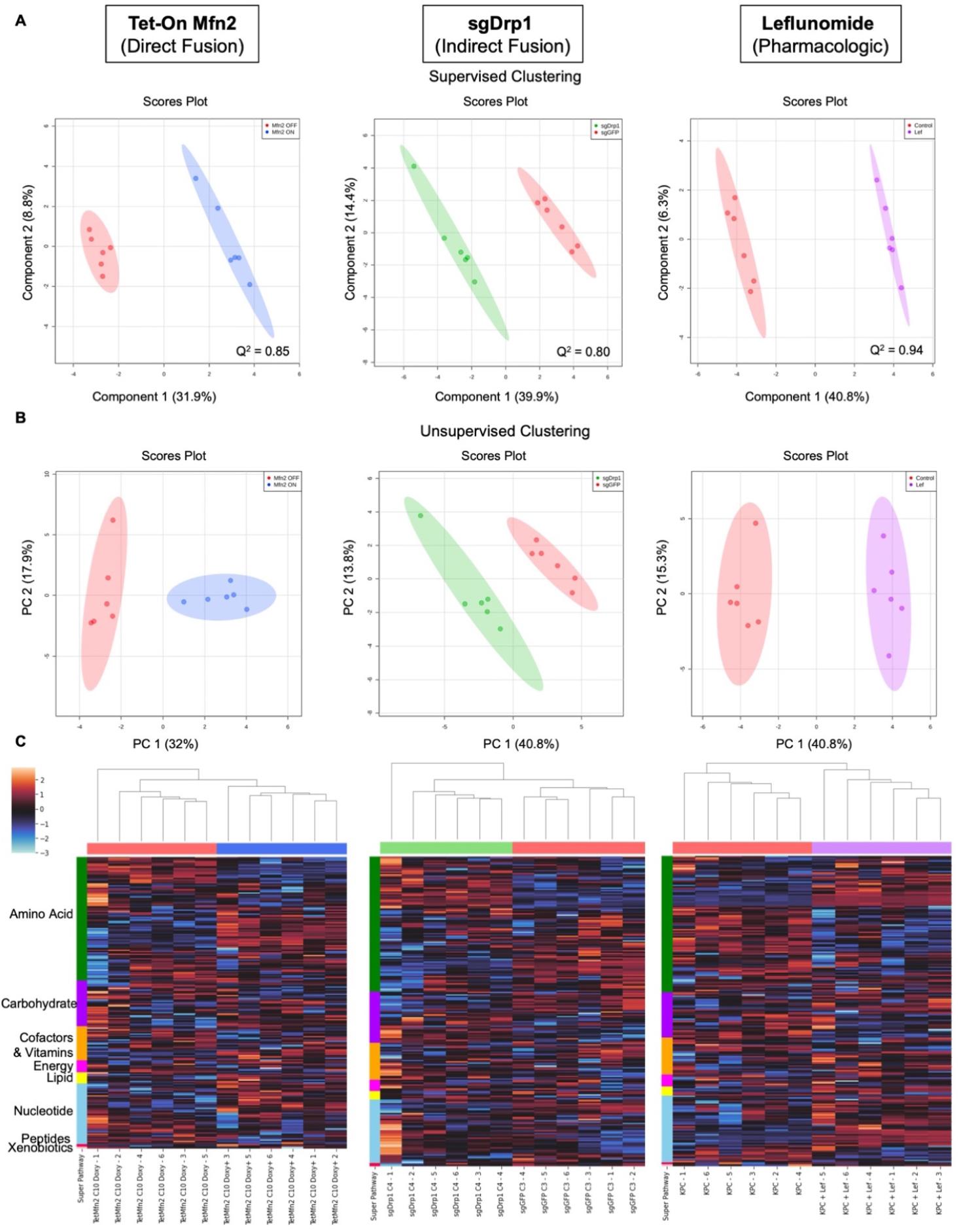
Multivariate clustering reveals distinct separation after inducing mitochondrial fusion when compared to controls. (A) Supervised PLS-DA and (B) unsupervised PCA score plots of Tet-On Mfn2 (blue), sgDrp1 (green), and Leflunomide (purple) treated KPC cells with respect to their corresponding controls (red). (C) Heatmap with unsupervised hierarchical clustering of affected super pathways across Tet-On Mfn2 (blue), sgDrp1 (green), and Leflunomide (purple). Both unsupervised and supervised clustering methods revealed a distinct separation between each method of fusion induction and its respective control. Predictive power of PLS-DA in component 1 represented by Q^2^ = 0.85 for Tet-On Mfn2, Q^2^ = 0.80 for sgDrp1, and Q^2^ = 0.94 for Leflunomide.

**Table 1.**
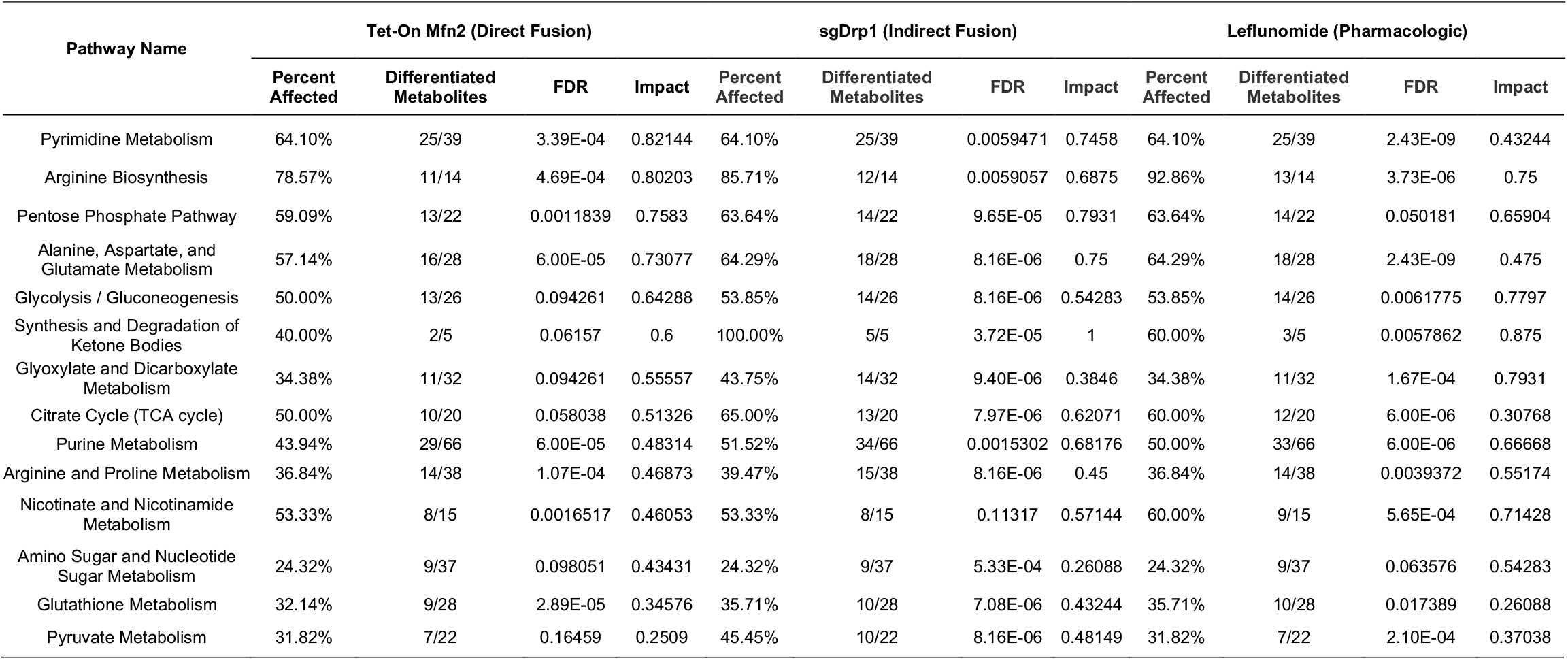
Common altered metabolic pathways from initial metabolite set across pathways supporting mitochondrial fusion. Pathway analysis included only pathways with an FDR < 0.05, impact > 0.25, and more than 20% of the metabolites in the pathway affected.

Notably, the following three Amino Acid pathways were significantly altered in the mitochondrial fusion cohorts compared to controls: arginine biosynthesis (FDR < 0.01, Impact > 0.68, and Percent Affected > 78.5%), alanine, aspartate, and glutamate metabolism (FDR < 0.0001, Impact > 0.47, and Percent Affected > 57.1%), and glutathione metabolism (FDR < 0.05, Impact > 0.26, and Percent Affected > 32.1%, Table 1). Pyrimidine and purine metabolism were the two nucleotide sub-pathways that were significantly altered after inducing mitochondrial fusion (FDR < 0.01, Impact > 0.43, Percent Affected = 64.1% and FDR < 0.01, Impact > 0.48, Percent Affected > 43.9% respectively, Table 1). We also observed significant changes in several carbohydrate metabolism sub-pathways, including the pentose phosphate pathway (PPP), glycolysis/gluconeogenesis, the citrate cycle (TCA cycle), pyruvate metabolism, and amino sugar and nucleotide sugar metabolism (Table 1). The full unfiltered pathway analysis for Tet-On Mfn2, sgDrp1, and Leflunomide treated KPC cells can be found in Figure 3 and Table S1.

**Figure 3.**
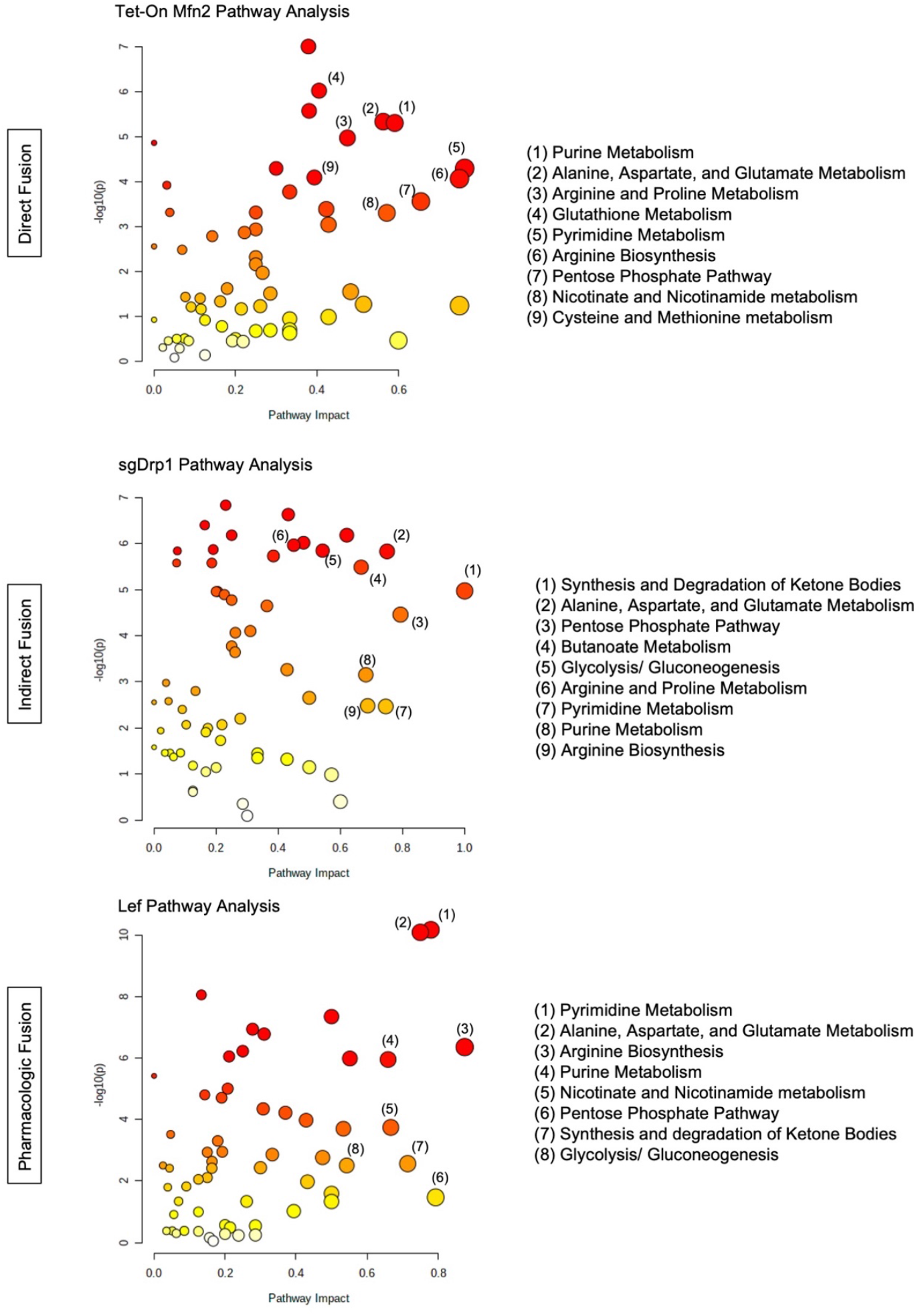
Total pathway analysis of filtered metabolites reveals similar impact of Amino Acid, Nucleotide, and Carbohydrate metabolism pathways as a function of mitochondrial fusion: Tet-On Mfn2 (direct), sgDrp1 (indirect), and leflunomide treatment (pharmacologic).

### Identification of Significantly Differentiated Metabolites

From this, we found that 75 out of 234 metabolites in Tet-On Mfn2, 54 out of 245 metabolites in the sgDrp1, and 74 out of 233 metabolites in the leflunomide treated groups were altered (both up-and-downregulated) compared to controls. As represented in their corresponding volcano plots, since LC-MS/MS was unable to detect many metabolites with a fold change greater than 2, we repeated our analysis with a lower stringency threshold, considering all significant metabolites based on an FDR < 0.05 in our initial univariate analysis (Figure 4A). A full list of discriminant metabolites identified via Student’s *t*-test can be found in Table S2A-C.

**Figure 4.**
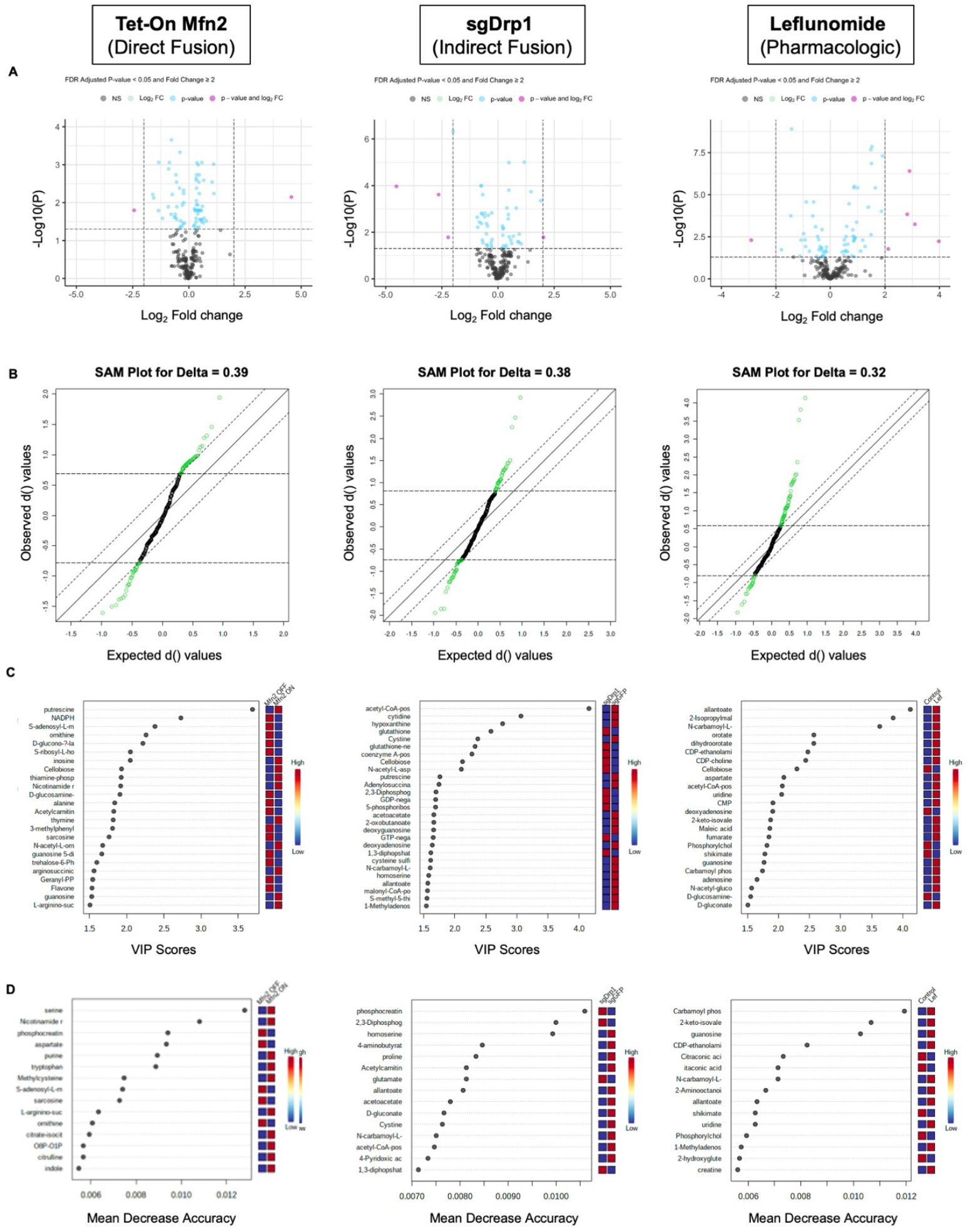
Statistical methods to identify differentially expressed metabolites after inducing mitochondrial fusion. (A) univariate Student’s *t*-test, FDR-adjusted *P*-value < 0.05, (B) SAM, FDR < 0.05, (C) PLS-DA, VIP score < 1.0, (D) RF model, Mean Decrease Accuracy > 0.

In order to ensure robustness in our feature selection process, we further developed three different pairwise models to identify metabolite markers for each group: a significance analysis of microarray (SAM) [11], PLS-DA variable importance in projection (VIP) [12], and a random forest (RF) [13] classification model. SAM identified 73 out of 234 metabolites in the Tet-On Mfn2 induced fusion group, 71 out of 245 metabolites in the indirect fusion group, and 74 out of 233 metabolites in the Leflunomide treated group as significantly altered based on an FDR < 0.05 and a corresponding delta of 0.39, 0.38, and 0.32 for the Tet-On Mfn2, sgDrp1, and Leflunomide groups respectively (Figure 4B). The full list of discriminant metabolites identified by SAM can be found in Table S3A-C.

Using our PLS-DA model, a VIP score greater than 1.0 [11] across all 5 principal components was used as a cutoff to identify discriminant metabolites after induction of mitochondrial fusion. From our original filtered metabolite set, we detected 83, 72, and 59 potential metabolites of interest in our Tet-On Mfn2, sgDrp1, and Leflunomide treated groups accordingly (Figure 4C and Table S4A-C). Moreover, permutation testing of 2000 repeats yielded a *P*-value = 0.001, suggesting that the separation exhibited by our PLS-DA model was not due to overfitting. We then performed leave-one-out cross validation [14] of the models and found that they had a predictive power 85% for Tet-On Mfn2, 80% for sgDrp1, and 94% for Leflunomide induced fusion.

To account for potential overfitting and potential bias from our previous models, we also developed an RF classification model for each group using MetaboAnalyst 5.0. For each RF model, we generated 500 trees to control for potential correlations between metabolites and subsequently measured a variable permutation importance score for each metabolite represented as the mean decrease accuracy (MDA) value. An MDA value approximates the amount that our model decreases in accuracy if the variable was taken out of the model [13]. Accordingly, we classified metabolites with MDA > 0 as discriminant and included them for pathway analysis. From our models, we identified 87 total discriminant metabolites in both the Tet-On Mfn2 and sgDrp1 groups and 81 total discriminant metabolites in the Leflunomide treated group (Figure 4D and Table S5A-C).

From our four statistical models, we combined the lists of significantly altered metabolites that contributed to each respective condition of induced mitochondrial fusion and only considered the overlap between all four lists for a definitive pathway analysis. This improved the robustness of our data analysis and further increased confidence in the identified metabolite markers for mitochondrial fusion in PDAC. As a result, we uncovered 48 unique identifier metabolites for direct fusion by Tet-On Mfn2 (Table 2), 38 unique identifier metabolites for indirect fusion by CRISPR knockout of Drp1 (Table 3), and 47 unique identifier metabolites for pharmacologic fusion by Leflunomide (Table 4).

**Table 2.**
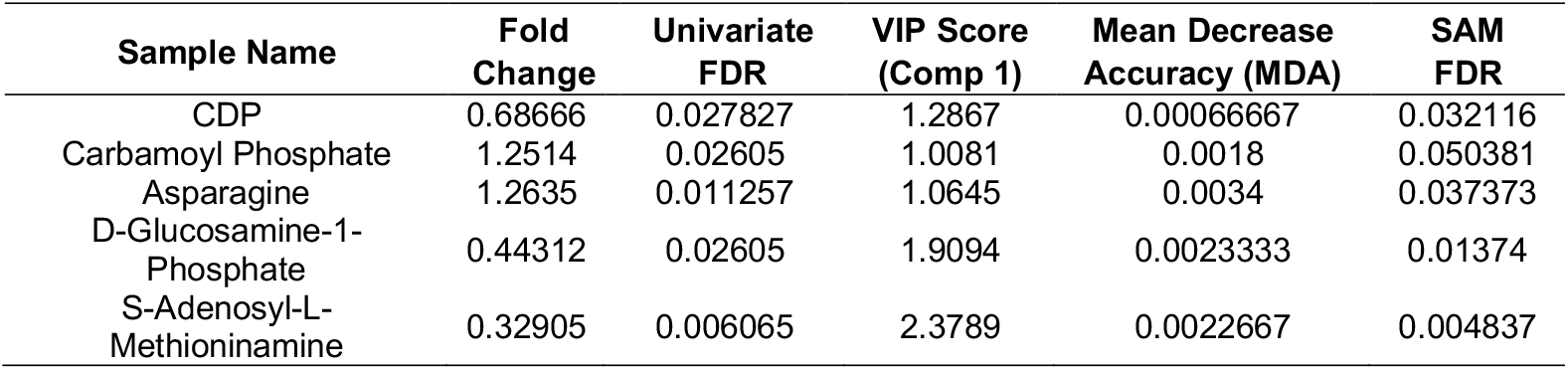
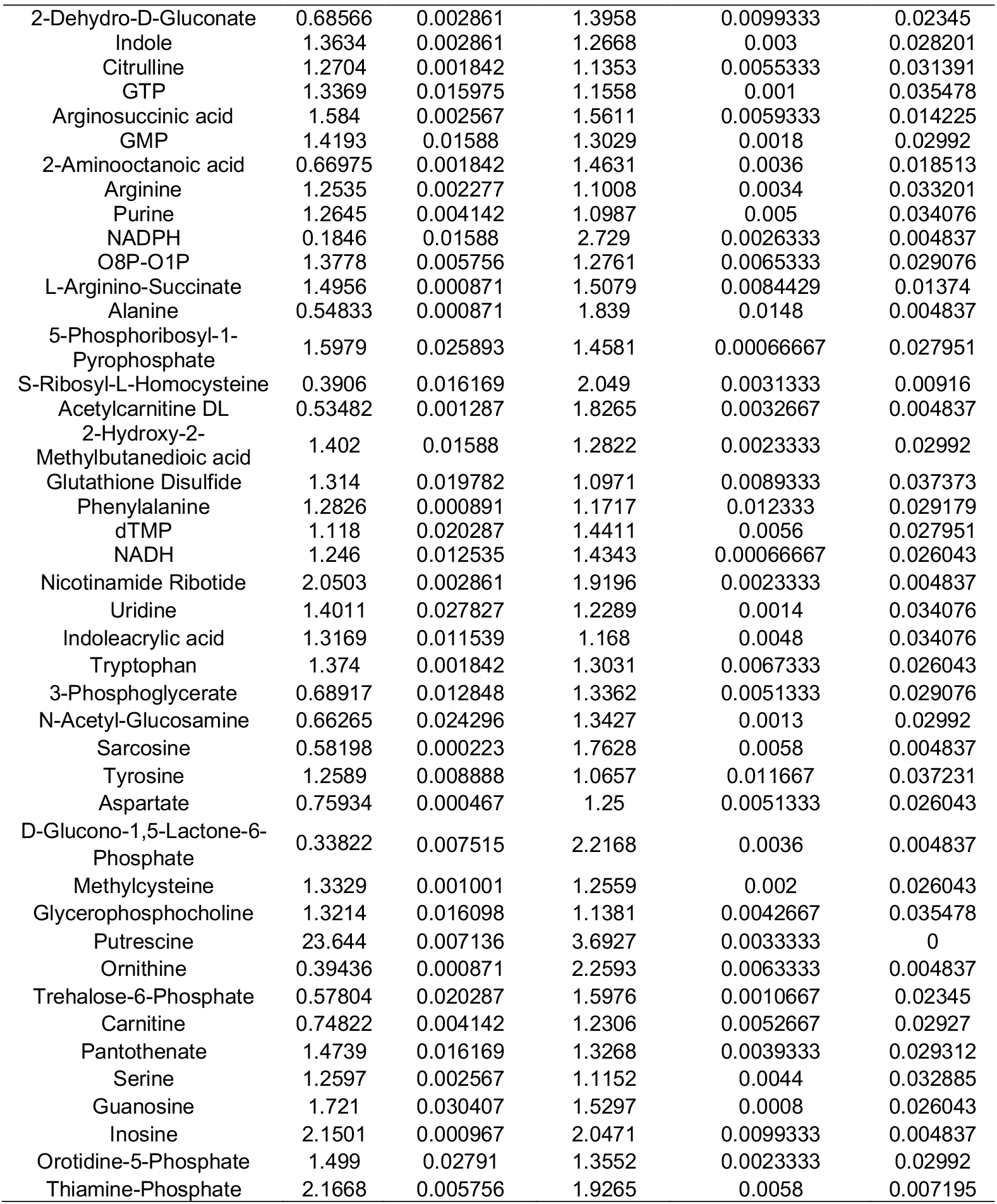
Discriminant metabolites identified after induction of Mfn2. Test statistics calculated for significantly altered metabolites overlapped across the univariate Student’s *t*-test, SAM, PLS-DA, and RF analysis.

| Sample Name | Fold Change | Univariate FDR | VIP Score (Comp 1) | Mean Decrease Accuracy (MDA) | SAM FDR |
| --- | --- | --- | --- | --- | --- |
| CDP | 0.68666 | 0.027827 | 1.2867 | 0.00066667 | 0.032116 |
| Carbamoyl Phosphate | 1.2514 | 0.02605 | 1.0081 | 0.0018 | 0.050381 |
| Asparagine | 1.2635 | 0.011257 | 1.0645 | 0.0034 | 0.037373 |
| D-Glucosamine-1-Phosphate | 0.44312 | 0.02605 | 1.9094 | 0.0023333 | 0.01374 |
| S-Adenosyl-L-Methioninamine | 0.32905 | 0.006065 | 2.3789 | 0.0022667 | 0.004837 |

Table 2. Cont.
| Sample Name | Fold Change | Univariate FDR | VIP Score (Comp 1) | Mean Decrease Accuracy (MDA) | SAM FDR |
| --- | --- | --- | --- | --- | --- |
| 2-Dehydro-D-Gluconate | 0.68566 | 0.002861 | 1.3958 | 0.0099333 | 0.02345 |
| Indole | 1.3634 | 0.002861 | 1.2668 | 0.003 | 0.028201 |
| Citrulline | 1.2704 | 0.001842 | 1.1353 | 0.0055333 | 0.031391 |
| GTP | 1.3369 | 0.015975 | 1.1558 | 0.001 | 0.035478 |
| Arginosuccinic acid | 1.584 | 0.002567 | 1.5611 | 0.0059333 | 0.014225 |
| GMP | 1.4193 | 0.01588 | 1.3029 | 0.0018 | 0.02992 |
| 2-Aminooctanoic acid | 0.66975 | 0.001842 | 1.4631 | 0.0036 | 0.018513 |
| Arginine | 1.2535 | 0.002277 | 1.1008 | 0.0034 | 0.033201 |
| Purine | 1.2645 | 0.004142 | 1.0987 | 0.005 | 0.034076 |
| NADPH | 0.1846 | 0.01588 | 2.729 | 0.0026333 | 0.004837 |
| O8P-O1P | 1.3778 | 0.005756 | 1.2761 | 0.0065333 | 0.029076 |
| L-Arginino-Succinate | 1.4956 | 0.000871 | 1.5079 | 0.0084429 | 0.01374 |
| Alanine | 0.54833 | 0.000871 | 1.839 | 0.0148 | 0.004837 |
| 5-Phosphoribosyl-1-Pyrophosphate | 1.5979 | 0.025893 | 1.4581 | 0.00066667 | 0.027951 |
| S-Ribosyl-L-Homocysteine | 0.3906 | 0.016169 | 2.049 | 0.0031333 | 0.00916 |
| Acetylcarnitine DL | 0.53482 | 0.001287 | 1.8265 | 0.0032667 | 0.004837 |
| 2-Hydroxy-2-Methylbutanedioic acid | 1.402 | 0.01588 | 1.2822 | 0.0023333 | 0.02992 |
| Glutathione Disulfide | 1.314 | 0.019782 | 1.0971 | 0.0089333 | 0.037373 |
| Phenylalanine | 1.2826 | 0.000891 | 1.1717 | 0.012333 | 0.029179 |
| dTMP | 1.118 | 0.020287 | 1.4411 | 0.0056 | 0.027951 |
| NADH | 1.246 | 0.012535 | 1.4343 | 0.00066667 | 0.026043 |
| Nicotinamide Ribotide | 2.0503 | 0.002861 | 1.9196 | 0.0023333 | 0.004837 |
| Uridine | 1.4011 | 0.027827 | 1.2289 | 0.0014 | 0.034076 |
| Indoleacrylic acid | 1.3169 | 0.011539 | 1.168 | 0.0048 | 0.034076 |
| Tryptophan | 1.374 | 0.001842 | 1.3031 | 0.0067333 | 0.026043 |
| 3-Phosphoglycerate | 0.68917 | 0.012848 | 1.3362 | 0.0051333 | 0.029076 |
| N-Acetyl-Glucosamine | 0.66265 | 0.024296 | 1.3427 | 0.0013 | 0.02992 |
| Sarcosine | 0.58198 | 0.000223 | 1.7628 | 0.0058 | 0.004837 |
| Tyrosine | 1.2589 | 0.008888 | 1.0657 | 0.011667 | 0.037231 |
| Aspartate | 0.75934 | 0.000467 | 1.25 | 0.0051333 | 0.026043 |
| D-Glucono-1,5-Lactone-6-Phosphate | 0.33822 | 0.007515 | 2.2168 | 0.0036 | 0.004837 |
| Methylcysteine | 1.3329 | 0.001001 | 1.2559 | 0.002 | 0.026043 |
| Glycerophosphocholine | 1.3214 | 0.016098 | 1.1381 | 0.0042667 | 0.035478 |
| Putrescine | 23.644 | 0.007136 | 3.6927 | 0.0033333 | 0 |
| Ornithine | 0.39436 | 0.000871 | 2.2593 | 0.0063333 | 0.004837 |
| Trehalose-6-Phosphate | 0.57804 | 0.020287 | 1.5976 | 0.0010667 | 0.02345 |
| Carnitine | 0.74822 | 0.004142 | 1.2306 | 0.0052667 | 0.02927 |
| Pantothenate | 1.4739 | 0.016169 | 1.3268 | 0.0039333 | 0.029312 |
| Serine | 1.2597 | 0.002567 | 1.1152 | 0.0044 | 0.032885 |
| Guanosine | 1.721 | 0.030407 | 1.5297 | 0.0008 | 0.026043 |
| Inosine | 2.1501 | 0.000967 | 2.0471 | 0.0099333 | 0.004837 |
| Orotidine-5-Phosphate | 1.499 | 0.02791 | 1.3552 | 0.0023333 | 0.02992 |
| Thiamine-Phosphate | 2.1668 | 0.005756 | 1.9265 | 0.0058 | 0.007195 |

**Table 3.** Discriminant metabolites identified after deletion of Drp1. Test statistics calculated for significantly altered metabolites overlapped across the univariate Student’s *t*-test, SAM, PLS-DA, and RF analysis.

| Sample Name | Fold Change | Univariate FDR | VIP Score (Comp 1) | Mean Decrease Accuracy (MDA) | SAM FDR |
| --- | --- | --- | --- | --- | --- |
| Betaine | 0.664180 | 0.002147 | 1.4133 | 0.0086 | 0.007697 |
| 4-Pyridoxic acid | 0.725180 | 0.037598 | 1.1296 | 0.0013333 | 0.033019 |
| Phosphocreatine | 1.316600 | 0.000906 | 1.1742 | 0.0052667 | 0.014145 |
| Aminoimidazole Carboxamide Ribonucleotide | 0.630100 | 0.028337 | 1.4028 | 0.0016667 | 0.014111 |
| Glutathione | 3.742600 | 0.000442 | 2.5826 | 0.0062667 | 0 |
| Glutathione | 3.742600 | 0.000182 | 2.3268 | 0.0049333 | 0 |
| Acetoacetate | 0.596270 | 0.000101 | 1.662 | 0.0072 | 0.004233 |
| 2-Oxobutanoate | 0.593130 | 0.000101 | 1.6618 | 0.0033333 | 0.004233 |
| GTP | 1.920100 | 0.026715 | 1.6525 | 0.0042667 | 0.007697 |
| N-Carbamoyl-L-Aspartate | 0.821900 | 0.001503 | 1.6041 | 0.0068 | 0.005064 |
| D-Gluconate | 0.724880 | 0.005425 | 1.2393 | 0.0022667 | 0.015196 |
| Homoserine | 0.598390 | 0.00421 | 1.5808 | 0.0054667 | 0.005681 |
| Acetyl-CoA | 0.043581 | 0.000107 | 4.1582 | 0.0039333 | 0 |
| N-Acetyl-L-Aspartic acid | 2.249000 | 9.87E-06 | 2.1057 | 0.0046667 | 0 |
| Adenylosuccinate | 0.533700 | 0.006098 | 1.7461 | 0.0057667 | 0.005064 |
| GDP | 2.053600 | 0.03156 | 1.6927 | 0.0043333 | 0.007697 |
| 5-Phosphoribosyl-1-Pyrophosphate | 1.758200 | 0.000906 | 1.6914 | 0.0022667 | 0.00449 |
| Cytidine | 0.159910 | 0.000242 | 3.0649 | 0.0071333 | 0 |
| S-Ribosyl-L-Homocysteine | 1.415700 | 0.015789 | 1.2606 | 0.00066667 | 0.016006 |
| Acetylcarnitine DL | 0.666230 | 0.001503 | 1.4281 | 0.0023333 | 0.007362 |
| N-Acetyl-Glutamine | 1.607500 | 0.014761 | 1.4623 | 0.0017333 | 0.01127 |
| Deoxyguanosine | 0.562700 | 0.003641 | 1.6583 | 0.0024667 | 0.005064 |
| Betaine Aldehyde | 0.702120 | 0.00421 | 1.3118 | 0.0048 | 0.012147 |
| 1,3-Diphosphoglycerate | 1.842500 | 0.025568 | 1.6151 | 0.0041333 | 0.00898 |
| Homocysteine | 1.377000 | 0.001738 | 1.2676 | 0.0057333 | 0.012147 |
| dAMP | 1.372600 | 0.002658 | 1.2376 | 0.0054 | 0.014111 |
| D-Glucono-1,5-Lactone-6-Phosphate | 0.700130 | 0.011761 | 1.2703 | 0.0018 | 0.015485 |
| Homocysteic acid | 0.616910 | 0.021676 | 1.4179 | 0.0014667 | 0.013178 |
| Cystine | 0.214900 | 0.016467 | 2.3705 | 0.0048 | 0.003848 |
| 4-Aminobutyrate | 0.741390 | 0.001858 | 1.2177 | 0.0078667 | 0.014111 |
| Putrescine | 0.528260 | 0.00226 | 1.7602 | 0.0076 | 0.00449 |
| Ornithine | 1.405100 | 1.04E-05 | 1.3596 | 0.0077667 | 0.005064 |
| coenzyme A | 4.070400 | 0.016467 | 2.2758 | 0.0041333 | 0.004233 |
| 2,3-Diphosphoglyceric acid | 1.926400 | 0.012063 | 1.6976 | 0.0048 | 0.00558 |
| Hypoxanthine | 0.251200 | 4.76E-07 | 2.7704 | 0.0075333 | 0 |

**Table 3. Cont.**
| Sample Name | Fold Change | Univariate FDR | VIP Score (Comp 1) | Mean Decrease Accuracy (MDA) | SAM FDR |
| --- | --- | --- | --- | --- | --- |
| Citrate | 1.398700 | 0.000156 | 1.329 | 0.0092667 | 0.007362 |
| Allantoate | 0.624710 | 0.000242 | 1.5666 | 0.001 | 0.004737 |
| 1-Methyladenosine | 0.614980 | 0.00169 | 1.5425 | 0.0023333 | 0.005064 |

**Table 4.** Discriminant metabolites identified after treatment with Leflunomide. Test statistics calculated for significantly altered metabolites overlapped across the univariate Student’s *t*-test, SAM, PLS-DA, and RF analysis.

| Sample Name | Fold Change | Univariate FDR | VIP Score (Comp 1) | Mean Decrease Accuracy (MDA) | SAM FDR |
| --- | --- | --- | --- | --- | --- |
| Citrate-Isocitrate | 0.653650 | 2.70E-05 | 1.1599 | 0.003967 | 0.007573 |
| CDP | 1.771000 | 0.001233 | 1.3036 | 0.005133 | 0.00732 |
| Carbamoyl Phosphate | 2.619400 | 5.46E-05 | 1.742 | 0.007 | 0.001114 |
| Fumarate | 2.787300 | 2.11E-08 | 1.8458 | 0.002514 | 0.000255 |
| Aminoimidazole Carboxamide Ribonucleotide | 0.526080 | 0.004786 | 1.3578 | 0.010638 | 0.007407 |
| Choline | 0.533840 | 0.011992 | 1.2772 | 0.002 | 0.008622 |
| Orotate | 15.739000 | 0.005825 | 2.5694 | 0.0008 | 0.000727 |
| Thiamine Pyrophosphate | 1.780700 | 0.002856 | 1.2673 | 0.003433 | 0.007573 |
| Acetoacetate | 0.586710 | 0.022826 | 1.1496 | 0.002267 | 0.015481 |
| Phosphorylcholine | 0.373280 | 1.28E-09 | 1.8143 | 0.011533 | 0.000255 |
| Isocitrate | 0.476540 | 0.00499 | 1.4156 | 0.002133 | 0.00732 |
| Deoxyadenosine | 0.288340 | 0.018626 | 1.9068 | 0.0009 | 0.005729 |
| 1-Methyladenosine | 1.883400 | 0.003598 | 1.36 | 0.006667 | 0.00732 |
| 2-Aminooctanoic acid | 1.811500 | 0.00499 | 1.2672 | 0.005743 | 0.007639 |
| D-Gluconate | 1.999000 | 3.90E-06 | 1.5065 | 0.005733 | 0.003395 |
| 2-Keto-Isovalerate | 2.868000 | 1.39E-08 | 1.8731 | 0.005743 | 0.000255 |
| Acetyl-CoA-Posi | 4.344300 | 0.016695 | 2.0614 | 0.0008 | 0.004201 |
| N-Carbamoyl-L-Aspartate | 51.968000 | 2.86E-10 | 3.627 | 0.0074 | 0 |
| Cellobiose | 0.133540 | 0.00499 | 2.2961 | 0.005467 | 0.001019 |
| O8P-O1P | 1.511200 | 0.000757 | 1.1186 | 0.008533 | 0.00911 |
| Thiamine-Phosphate | 1.822400 | 0.0487 | 1.1472 | 0.0005 | 0.018784 |
| Creatine | 1.813900 | 3.86E-06 | 1.394 | 0.004333 | 0.00544 |
| CDP-Ethanolamine | 7.034600 | 0.000145 | 2.4725 | 0.005 | 0 |
| Acetylcarnitine DL | 1.841000 | 0.00338 | 1.2953 | 0.004133 | 0.007573 |
| Aconitate | 0.542060 | 2.70E-05 | 1.3988 | 0.009033 | 0.005729 |
| Shikimate | 0.366580 | 0.000181 | 1.7807 | 0.005267 | 0.001198 |
| Anthranilate | 1.485100 | 0.021202 | 1.0136 | 0.002333 | 0.021203 |

Table 4. Cont.
| Sample Name | Fold Change | Univariate FDR | VIP Score (Comp 1) | Mean Decrease Accuracy (MDA) | SAM FDR |
| --- | --- | --- | --- | --- | --- |
| Uridine | 3.660400 | 9.62E-05 | 2.0507 | 0.001143 | 0.000727 |
| 2-Isopropylmalic acid | 81.792000 | 1.87E-11 | 3.8439 | 0.009533 | 0 |
| CMP | 3.126500 | 3.90E-06 | 1.9097 | 0.004067 | 0.000424 |
| CDP-Choline | 8.569100 | 0.000571 | 2.4383 | 0.0036 | 0.000255 |
| Deoxyguanosine | 0.507480 | 0.002085 | 1.4019 | 0.0026 | 0.00732 |
| Citraconic acid | 0.639440 | 0.000181 | 1.1726 | 0.004733 | 0.007573 |
| N-Acetyl-Glucosamine | 1.556400 | 0.008589 | 1.109 | 0.005467 | 0.01403 |
| Glycerophosphocholine | 1.606000 | 3.48E-05 | 1.2246 | 0.004 | 0.00732 |
| 2-Oxo-4-Methylthiobutanoate | 0.461890 | 0.043799 | 1.4045 | 0.0014 | 0.00941 |
| Histidinol | 1.546000 | 0.024378 | 1.021 | 0.0032 | 0.021203 |
| 4-Aminobutyrate | 1.947500 | 0.000426 | 1.4171 | 0.0046 | 0.006111 |
| Dihydroorotate | 7.470800 | 3.97E-07 | 2.5685 | 0.0076 | 0 |
| UDP | 1.853800 | 0.004218 | 1.3149 | 0.0066 | 0.007573 |
| Itaconic acid | 0.688280 | 0.000804 | 1.0502 | 0.006067 | 0.012297 |
| Maleic acid | 2.836300 | 1.41E-07 | 1.8604 | 0.0048 | 0.000424 |
| dCDP | 1.193600 | 0.00578 | 1.3101 | 0.004667 | 0.007639 |
| N-Acetyl-Glucosamine-1-Phosphate | 2.279300 | 0.00499 | 1.5674 | 0.0008 | 0.006463 |
| Aspartate | 3.759100 | 4.98E-08 | 2.0869 | 0.0108 | 0 |
| Allantoate | 158.36 | 3.66E-12 | 4.1202 | 0.0084 | 0 |
| Guanosine | 2.803700 | 0.002386 | 1.7677 | 0.005 | 0.003886 |

### Targeted Pathway Analysis Distinguishes altered Metabolome after Mitochondrial Fusion

When conducting sub-pathway analysis from each list of discriminant metabolites identified via one of our four statistical models, we found that the overarching patterns observed in alterations of Amino Acid, Nucleotide, and Carbohydrate super-pathways remained similar to those of our initial untargeted sub-pathway analysis. More importantly, sub-pathway analysis from each distinct discriminant metabolite identification method appeared to yield very similar phenotypes across direct genetic fusion, indirect genetic fusion, and pharmacologic fusion (Figures S1-4 and Tables S6-9A-C). Using our overlapped discriminant metabolite list for sub-pathway analysis, we discovered that even our more limited metabolite set was able to recapitulate these trends in metabolic repogramming in each independent method of mitochondrial fusion induction. Specifically, Amino Acid, Nucleotide, and Carbohydrate pathways were altered across all three experimental groups (Figure 5). We found eight particular sub-pathways that were considered significantly impacted after filtering the raw pathway outputs from MetaboAnalyst based on an FDR < 0.05 (Table S10A-C). These included alanine, aspartate, and glutamate metabolism (FDR < 0.0001), arginine biosynthesis (FDR < 0.0001), glutathione metabolism (FDR < 0.05), cysteine and methionine metabolism (FDR < 0.01), pyrimidine metabolism (FDR < 0.0001), purine metabolism (FDR < 0.0001), PPP (FDR < 0.001), and glycolysis/gluconeogenesis (FDR < 0.05, Table 5).

**Figure 5.**
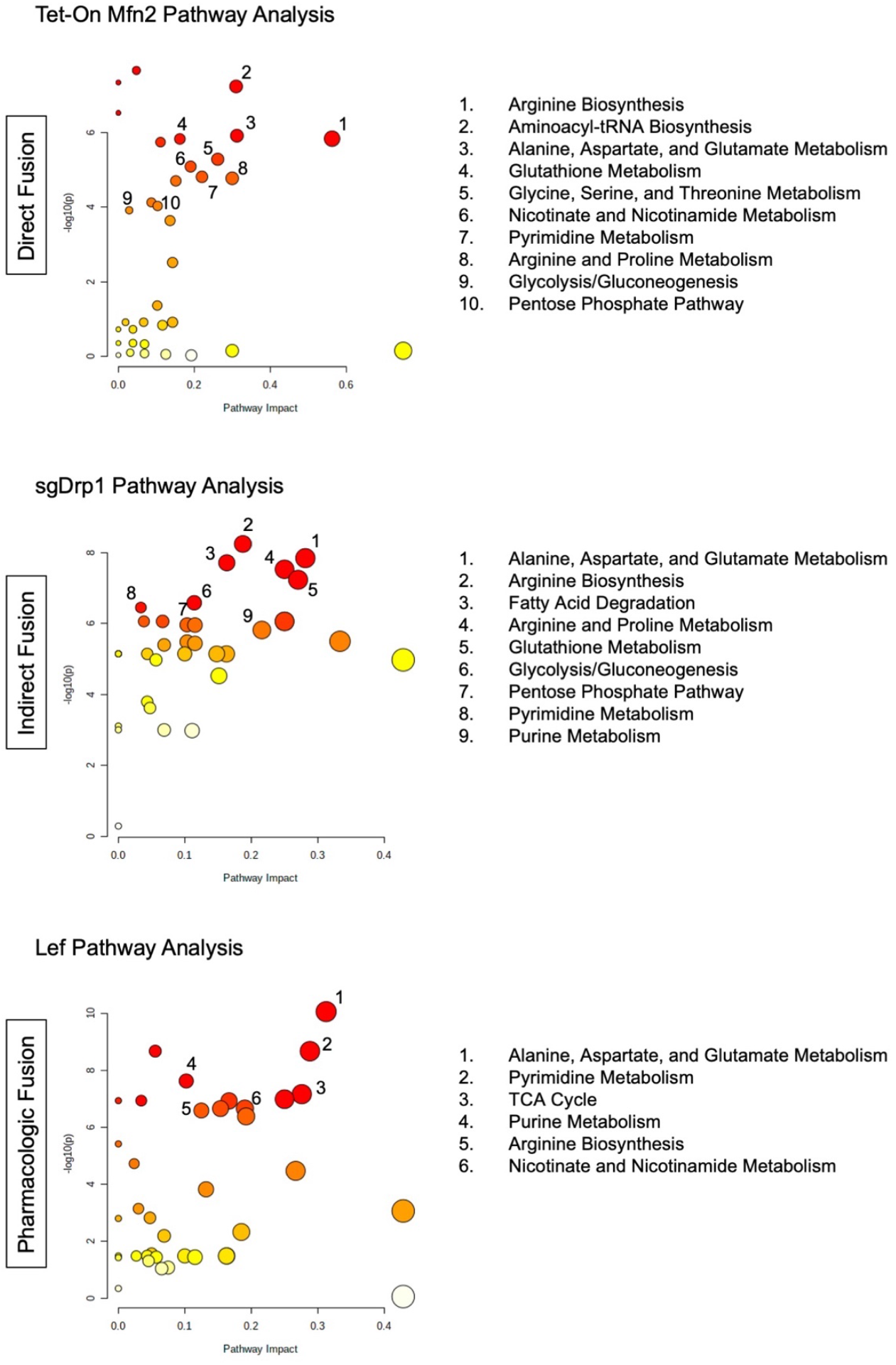
Pathway analysis of overlapped discriminant metabolites from direct fusion in Tet-On Mfn2, indirect fusion in sgDrp1, and pharmacologic fusion in leflunomide treated KPC cells.

**Table 5.** Significantly altered pathways from overlapped discriminant metabolite sets.

| Pathway Name | Percent Affected | Differentiated Metabolites | FDR | Impact |
| --- | --- | --- | --- | --- |
| *Tet-On Mfn2 (Direct Fusion)* |  |  |  |  |
| Beta-Alanine Metabolism | 9.52% | 2/21 | 6.74E-07 | 0.04762 |
| Aminoacyl-tRNA Biosynthesis | 16.67% | 8/48 | 6.74E-07 | 0.31033 |

Table 5. Cont.
| Pathway Name | Percent Affected | Differentiated Metabolites | FDR | Impact |
| --- | --- | --- | --- | --- |
| *Tet-On Mfn2 (Direct Fusion)* |  |  |  |  |
| Alanine, Aspartate, and Glutamate Metabolism* | 28.57% | 8/28 | 7.36E-06 | 0.3125 |
| Arginine Biosynthesis* | 42.86% | 6/14 | 7.36E-06 | 0.5625 |
| Glutathione Metabolism* | 14.29% | 4/28 | 7.36E-06 | 0.16216 |
| Pantothenate and CoA Biosynthesis | 15.79% | 3/19 | 7.85E-06 | 0.11112 |
| Glycine, Serine, and Threonine Metabolism | 17.65% | 6/34 | 0.00002 | 0.26191 |
| Nicotinate and Nicotinamide Metabolism | 13.33% | 2/15 | 2.83E-05 | 0.19048 |
| Pyrimidine Metabolism* | 23.08% | 9/39 | 4.87E-05 | 0.22035 |
| Arginine and Proline Metabolism | 13.16% | 5/38 | 4.88E-05 | 0.3 |
| Cysteine and Methionine Metabolism* | 15.15% | 5/33 | 5.28E-05 | 0.15151 |
| Amino Sugar and Nucleotide Sugar Metabolism | 5.41% | 2/37 | 0.000185 | 0.08696 |
| Pentose Phosphate Pathway* | 13.64% | 3/22 | 0.000216 | 0.10345 |
| Glycolysis / Gluconeogenesis* | 3.85% | 1/26 | 0.000264 | 0.02857 |
| Purine Metabolism* | 10.61% | 7/66 | 0.00047 | 0.13634 |
| *sgDrp1 (Indirect Fusion)* |  |  |  |  |
| Arginine Biosynthesis* | 14.29% | 2/14 | 2.05E-07 | 0.1875 |
| Alanine, Aspartate, and Glutamate Metabolism* | 21.43% | 6/28 | 2.32E-07 | 0.28125 |
| Fatty Acid Degradation | 5.13% | 2/39 | 2.32E-07 | 0.16327 |
| Arginine and Proline Metabolism | 15.79% | 6/38 | 2.66E-07 | 0.25 |
| Glutathione Metabolism* | 17.86% | 5/28 | 4.20E-07 | 0.27028 |
| Glycolysis / Gluconeogenesis* | 11.54% | 3/26 | 1.56E-06 | 0.11428 |
| Pyrimidine Metabolism* | 7.69% | 3/39 | 1.82E-06 | 0.0339 |
| Nitrogen Metabolism | 16.67% | 1/6 | 2.85E-06 | 0.25 |
| Pentose Phosphate Pathway* | 18.18% | 4/22 | 3.06E-06 | 0.10344 |
| Glyoxylate and Dicarboxylate Metabolism | 9.38% | 3/32 | 3.06E-06 | 0.11538 |
| Purine Metabolism* | 15.15% | 10/66 | 3.92E-06 | 0.2159 |
| Butanoate Metabolism | 26.67% | 4/15 | 7.42E-06 | 0.33334 |

Table 5. Cont.
| ***sgDrp1 (Indirect Fusion)*** |  |  |  |  |
| --- | --- | --- | --- | --- |
| Citrate Cycle (TCA cycle) | 10.00% | 2/20 | 7.42E-06 | 0.10345 |
| Propanoate Metabolism | 8.70% | 2/23 | 7.69E-06 | 0.11538 |
| Cysteine and Methionine Metabolism* | 12.12% | 4/33 | 3.69E-05 | 0.15151 |
| ***Leflunomide (Pharmacologic)*** |  |  |  |  |
| Alanine, Aspartate, and Glutamate Metabolism* | 25.00% | 7/28 | 3.28E-09 | 0.3125 |
| Pantothenate and CoA Biosynthesis | 10.53% | 2/19 | 2.69E-08 | 0.05556 |
| Pyrimidine Metabolism* | 25.64% | 10/39 | 2.69E-08 | 0.28815 |
| Purine Metabolism* | 9.09% | 6/66 | 2.25E-07 | 0.10227 |
| Citrate Cycle (TCA cycle) | 30.00% | 6/20 | 5.03E-07 | 0.27587 |
| Valine, Leucine, and Isoleucine Biosynthesis | 12.50% | 1/8 | 5.03E-07 | 0.25 |
| Aminoacyl-tRNA Biosynthesis | 2.08% | 1/48 | 5.03E-07 | 0.03448 |
| Glyoxylate and Dicarboxylate Metabolism | 12.50% | 4/32 | 7.65E-07 | 0.15384 |
| Nicotinate and Nicotinamide Metabolism | 13.33% | 2/15 | 7.65E-07 | 0.19048 |
| Arginine Biosynthesis* | 21.43% | 3/14 | 8.20E-07 | 0.125 |
| Glycine, Serine, and Threonine Metabolism | 5.88% | 2/34 | 4.87E-05 | 0.02381 |
| Butanoate Metabolism | 26.67% | 4/15 | 8.19E-05 | 0.26667 |
| Valine, Leucine, and Isoleucine Degradation | 10.00% | 4/40 | 0.000343 | 0.13208 |
| Cysteine and Methionine Metabolism* | 3.03% | 1/33 | 0.00155 | 0.0303 |
| Beta-Alanine Metabolism | 9.52% | 2/21 | 0.002936 | 0.04762 |
| Pentose Phosphate Pathway* | 9.09% | 2/22 | 0.010995 | 0.06896 |
| Glutathione Metabolism* | 3.57% | 1/28 | 0.04232 | 0.02703 |
| Glycolysis / Gluconeogenesis* | 7.69% | 2/26 | 0.043741 | 0.05714 |

\*Pathways common amongst Tet-On Mfn2, sgDrp1, and Lef treatment.

Further analysis revealed that although these pathways were considered statistically significant based on our FDR adjusted *P*-value, we observed that many of these pathways had a low impact and correspondingly low percentage of affected metabolites in one of the three groups. Diving deeper, arginine biosynthesis appeared to be more affected after direct fusion with Tet-On Mfn2 (Impact = 0.56, Percent Affected = 42.9%) than after indirect fusion with sgDrp1 (Impact = 0.19, Percent Affected = 14.3%) and pharmacologic fusion with Leflunomide (Impact = 0.13, Percent Affected = 21.4% Table 5). Glutathione metabolism was more heavily affected by indirect fusion (Impact = 0.27, Percent Affected = 17.9%) than direct fusion (Impact = 0.16, Percent Affected = 14.3%) and pharmacologic fusion (Impact = 0.03, Percent Affected = 3.6%). Cysteine and methionine metabolism was least impacted by Leflunomide treatment (Impact = 0.03, Percent Affected = 3.0%) followed by Drp1 knockout (Impact = 0.15, Percent Affected = 12.1%) and Mfn2 upregulation (Impact = 0.15, Percent Affected = 15.2%, Table 5). Understandably, pyrimidine metabolism was most affected by Leflunomide treatment (Impact = 0.28, Percent Affected = 25.6%) since its mechanism of action directly inhibits dihydroorotate dehydrogenase (DHODH), a crucial enzyme in the de novo pyrimidine biosynthesis pathway. Interestly, direct fusion through Tet-On Mfn2 closely mirrored Leflunomide’s effect on pyrimidine metabolism, having an impact of 0.22 on the pathway with 23.1% of its metabolites altered (Table 5). Purine metabolism was most affected by indirectly inducing fusion with knockout of Drp1 (Impact = 0.22, Percent Affected = 15.2%); though, still moderately altered via direct fusion with Tet-On Mfn2 (Impact = 0.14, Percent Affected = 10.6%) and pharmacologic fusion with Leflunomide (Impact = 0.10, Percent Affected = 9.1%, Table 5). Both carbohydrate metabolism sub-pathways, PPP, and glycolysis were modestly impacted across the three groups, but knockdown of Drp1 in particular had more impact on glycolysis/gluconeogenesis (Impact = 0.11, Percent Affected = 11.5%), supporting recent findings that Drp1 promotes metabolic changes through glycolysis to drive PDAC tumorigenesis (Table 5) [7,15]. The main pathway that had an impact greater than 0.28 across the Tet-On Mfn2, sgDrp1, and Leflunomide groups was alanine, aspartate, and glutamate metabolism with more than 21.4% of the pathway appearing significantly altered (Table 5).

Interestingly, several pathways from this analysis were identified as specific to each independent method for mitochondrial fusion induction. We noticed that direct fusion via Tet-On Mfn2 showed distinct impact on aminoacyl-tRNA biosynthesis and glycine, serine, and threonine metabolism in the top 5 pathway hits (Figure 5). Likewise, fatty acid degradataion and the citrate cycle were specific to sgDrp1 and Leflunomide groups when considering only the top 5 pathway hits (Figure 5). Nevertheless, after mapping the significantly altered metabolic pathways identified from the KEGG database using our overlapped discriminant metabolite set exhibited that they were in fact highly interconnected (Figure S5). Alanine, aspartate, and glutamate metabolism fed into each of the previously mentioned metabolic pathways, aligning with each of our metabolic screens. Furthermore, we see that many of the pathways are interdependent among each other, suggesting that altering mitochondrial morphology from a punctate to fused state does in fact play a significant role in metabolic reprogramming in favor of curbing tumorigenesis.

## Discussion

Advances in mitochondrial biology in the previous decade have opened the doors to novel means of therapeutically targeting tumorigenesis. It has been widely shown that mitochondrial respiration is essential across multiple tumor-types in order to circumvent limitations in glycolysis, actively remodeling their means for cellular energetics [3,16–18]. This is particularly true in pancreatic cancer where mitochondrial dysfunction has been found to shift the cellular bioenergetics of cells to favor OXPHOS, supporting proliferation and metastasis [19,20]. Moreover, we and others have shown that defects in *KRAS,* the most widely mutated gene in PDAC, is characteristic of fragmented mitochondria [1,7,20]. Although more than 90% of PDAC cases are driven by oncogenic *KRAS* [21], limited advancements have been made in formulating a clinical approach to target the gene. Instead, our work provided an alternative solution, altering the phenotypic state of mitochondria driven by KRAS through Drp1, which in turn suppressed tumor growth through modulating levels of defective mitochondria and limiting OXPHOS capability in PDAC [1]. In order to gain a better understanding of the impact of shifting the status of mitochondria from fragmented to fused, we attempted to elucidate the metabolome of PDAC after three independent methods of inducing fusion. To our knowledge, this is the first comparative metabolomic study of mitochondrial fusion in pancreatic cancer to identify macroscopic pathway alterations.

Our study builds upon previous findings to better understand common metabolic perturbations from mitochondrial fusion. Here, we used three different models: a direct fusion model involving the upregulation of Mfn2 in a doxycycline-dependent manner, an indirect fusion model through the attenuation of Drp1 using CRISPR-Cas9, and a pharmacologic approach with Leflunomide, each in KPC cells. Of note, alanine, aspartate, and glutamate metabolism was the most prominent hit across all three methods of induced fusion, understandably because of its overarching position providing the precursors for the other seven identified metabolic pathways commonly altered (Figures 5 and S5). This finding further supports current working theories of tumor ability to alternatively fuel metabolism using extracellular amino acid pools, particularly, alanine, aspartate, glutamate, and asparagine, as carbon sources [10,22].

We also observed several expected alterations in nucleotide pathways downstream due to dysregulation of glycolysis and PPP that are linked to mutant KRAS activation [20,23–25] by reducing the pool of fragmented mitochondria present. Leflunomide is widely known for its direct inhibition of DHODH, an enzyme localized on the inner mitochondrial membrane, that is responsible for de novo pyrimidine biosynthesis. Interestingly, genetic modulation of mitochondrial morphology appeared to affect these pathways in a similar fashion. Tet-On Mfn2 and sgDrp1 modulated pyrimidine biosynthesis in addition to glycolysis and PPP, showing a common set of metabolic disturbances across multiple super-pathways. Ultimately, this suggests that mitochondrial fusion may actually be working in tandem with DHODH inhibition as a tumor suppressive mechanism in pancreatic cancer.

Nevertheless, future research is needed in order to fully characterize the mechanism by which mitochondrial fusion reprograms the metabolome to curb tumorigenesis. We recognize that our study follows a markedly stringent method of discriminant metabolite identification. Relative metabolite concentration readings missed at least 1 data point for more than 20% of our metabolites across all three induced fusion groups as a result of either low concentration or poor mass spectrometry signal response, potentially limiting our analysis. Methods to impute these missing values should be explored when doing an in-depth analysis of sub-pathway alterations. However, our study provides foundational evidence that mitochondrial morphology plays a notable role in metabolic reprogramming further supporting leflunomide as a novel therapeutic against PDAC due to its ability to leverage both mitochondrial fusion and DHODH inhibition.

## Materials and Methods

### Cell Culture

Murine KPC cells syngeneic with C57BL/6 (K8484) were a generous gift from Anirban Maitra from the University of Texas MD Anderson Cancer Center. KPC cells were grown in RPMI-1640 supplemented with 10% FBS, 2 mM GlutaMax, 1 mM sodium pyruvate, and 7 μg/mL of insulin. We previously described the generation and selection of KPC Tet-On Mfn2, sgDrp1, and sgGFP clones [1].

### Untargeted Metabolomic Analysis

All metabolomic analyses were conducted under steady state conditions. KPC cell lines were grown in appropriate growth media in six replicates in 10-cm plates. Two hours before metabolite collection, cells were incubated in fresh growth media. Accordingly, replicate cell lines were plated and grown in parallel in order to control for cell growth. Cell counts from the replicate plates were used to normalize metabolite readings. After incubation in fresh growth media, 4 mL of 80% methanol that was pre-chilled to −80°C was added, and the cell plates were immediately transferred to −80°C to incubate overnight. The cell lysate-methanol mixture was then scraped and transferred to conical tubes on dry ice and centrifuged at 5000 g for 5 minutes. The supernatant was collected, and the process was repeated two more times after resuspending the pellet in 500 μL of chilled 80% methanol, for a total volume of 5 mL. Samples were then completely dried via speed vacuum at 30°C. Metabolites were analyzed by LC-MS/MS as previously described [9,10].

### Discriminant Metabolite Identification

From our initial 296 measured metabolites, we stringently filtered out readings that were missing any values to ensure robustness in our analysis. As a result, 79.1%, 82.8%, and 78.7% of the metabolites within the datasets remained for our Tet-On Mfn2, sgDrp1, and Leflunomide groups, respectively. Continued filtering for altered metabolites was performed using four independent statistical approaches: (1) FDR-adjusted two-sided Student’s *t*-test (*P*-values < 0.05 were considered statistically significant), (2) SAM (significant metabolite features were identified at an FDR < 0.05 and corresponding delta of 0.39 for Tet-On Mfn2, 0.38 for sgDrp1, and 0.32 for Leflunomide), (3) PLS-DA variable importance in projection (VIP scores > 1.0 were considered significant for class separation), and (4) RF classification based on 500 trees (significance established at permutation importance, MDA > 0).

Two-sided Student’s *t*-test characterized significant differences based on hypothesis testing of the groups’ means following a normal distribution. However, since we only had an n = 6 for each group, there is the possibility that the variance in our dataset is not stable [26]. To account for this, SAM uses a nonparametric approach which does not rely on a prescribed probability distribution [27,28]. SAM processes multiple permutations of our data in order to calculate FDR values, which we are able to control using the tuning parameter delta, allowing us to define our cutoff for identification of altered metabolites [29]. PLS-DA VIP measures the importance of each variable after supervised dimensional reduction using a partial least squares projection [30,31]. Given that the average squared VIP score is 1.0, we followed the field standard of considering VIP scores greater than 1.0 as significantly altered and confirmed its predictive probabilities using a leave-one-out cross validation method [32]. RF is a machine learning model often used for regression and classification. We tuned an RF model using bootstrap sampling to generate 500 random classification trees. Since this model only uses a subset of the available data to generate trees, we were able to robustly limit overfitting as well as potential outliers [33]. An MDA score was calculated using the unbiased out-of-bag classification error for each metabolite predictor, representing its predictive importance for the model. The reference MDA of 0 signifies that the predictor has no predictive importance in the model. Therefore, a metabolite with an MDA > 0 represents that the loss of that metabolite from the model will result in a decrease in predictive ability of the RF model, which we used as our cutoff for identifying discriminant metabolites. In order to confirm our pathway analysis findings from the discriminant metabolite lists generated from each of our statistical models, we further refined our list of metabolites by taking only those that were common across all four feature selection methods.

### Pathway Analysis

Custom data mining using BioPython’s KEGG API was used to collect super metabolic pathway data for hierarchical clustering similar to what has previously been described [34]. Further sub-pathway analysis was performed using MetaboAnalyst 5.0, initially on the total filtered metabolite set from the KEGG database for *mus musculus.* We used a Global Test for enrichment and out-degree centrality for topology analysis. Using the hit values from the MetaboAnalyst output, we calculated a percent affected score for each pathway to prevent identification of significantly altered pathways with fewer than 20% of their metabolites altered. Significant sub-pathways were also filtered using an FDR adjusted *P*-value < 0.05 and impact greater than 0.25. Confirmation of sub-pathway alterations after induction of mitochondrial fusion was performed after running continued pathway analyses of each discriminant metabolite set generated from our four different statistical methods as well as our overlapped metabolite dataset.

### Statistical Analysis

Data manipulation and statistical analyses were performed using MetaboAnalyst 5.0 [35], R version 4.0.2, and the Pandas, NumPy, and SciPy libraries in python 3.8. Metabolite concentration values were normalized based on the control group for each respective experimental group and log-transformed and pareto-scaled to approximate a normal distribution. 2D PCA and PLS-DA scores plots were generated using MetaboAnalyst. Heatmaps were generated with the python Seaborn package using a euclidean distance measure and ward algorithm. Volcano plots were generated using the EnhancedVolcano package from Bioconductor in R [36].

## Author Contributions

Conceptualization, N.D.N., M.Y. and C.M.T.; methodology, N.D.N. and M.Y.; validation, N.D.N., formal analysis, N.D.N.; investigation, N.D.N. and M.Y., resources, N.D.N. and M.Y.; data curation, N.D.N., M.Y., M.Y. and J.M.A.; writing —original draft preparation, N.D.N.; writing —review and editing, N.D.N., V.Y.R., E.C.M., J.A.J., and C.M.T.; visualization, N.D.N., V.Y.R., A.R., E.C.M., and J.A.J.; supervision, N.D.N., M.Y. and C.M.T.; project administration, N.D.N. and C.M.T., funding acquisition, C.M.T. All authors have read and agreed to the published version of the manuscript.

## Acknowledgments

Graphical abstract created with BioRender.com adapted from the “Untargeted Metabolomics for Discover of Disease Biomarkers” template by Anne-Laure Agrinier (2021) retrieve from https://app.biorender.com/biorender-templates.

## Data & Resource Sharing

Data, materials, and reagents will be made available to other researchers upon request.

