## Supplementary Figures 1-5 and Tables 1-10 for "Comparative untargeted metabolomic profiling of induced mitochondrial fusion in pancreatic cancer": Supplementary Figures 1-5.pdf

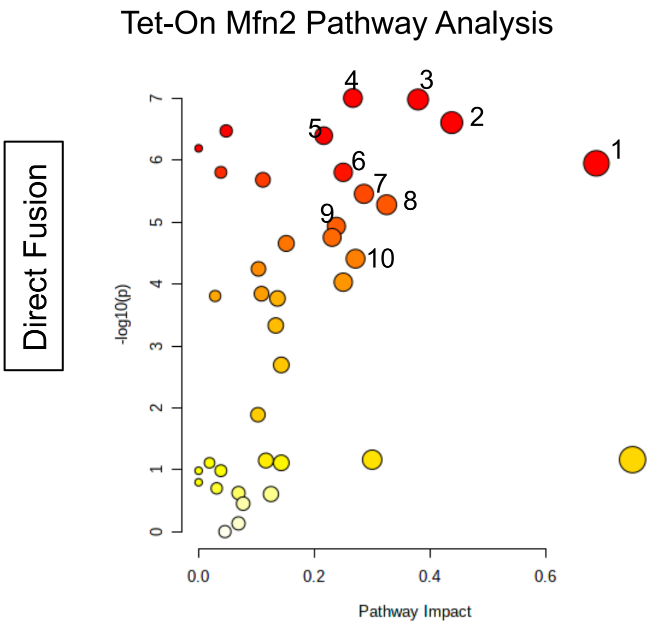


1. Arginine Biosynthesis
2. Alanine, Aspartate, and Glutamate Metabolism
3. Aminoacyl-tRNA Biosynthesis
4. Histidine Metabolism
5. Glutathione Metabolism
6. D-Glutamine and D-Glutamate Metabolism
7. Glycine, Serine, and Threonine Metabolism
8. Arginine and Proline Metabolism
9. Nicotinate and Nicotinamide Metabolism
10. Pyrimidine Metabolism

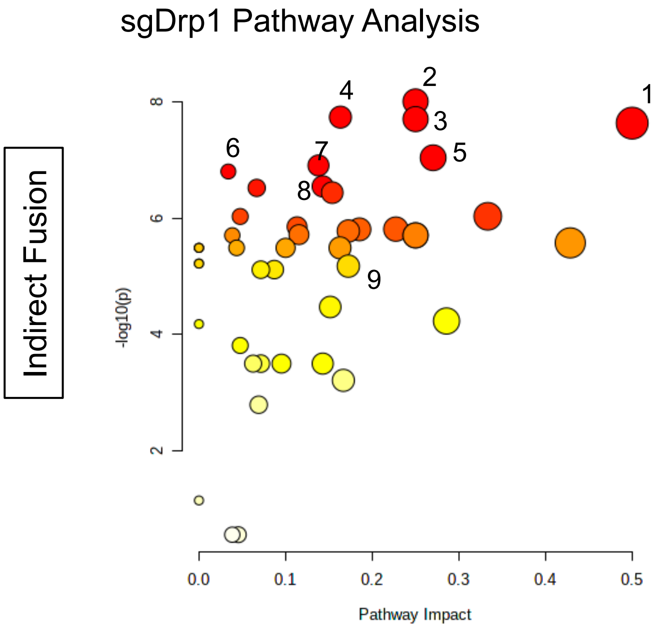


1. Alanine, Aspartate, and Glutamate Metabolism
2. Arginine and Proline Metabolism
3. Arginine Biosynthesis
4. Fatty Acid Degradation
5. Glutathione Metabolism
6. Pyrimidine Metabolism
7. Aminoacyl-tRNA Biosynthesis
8. Glycolysis/Gluconeogenesis
9. Pentose Phosphate Pathway

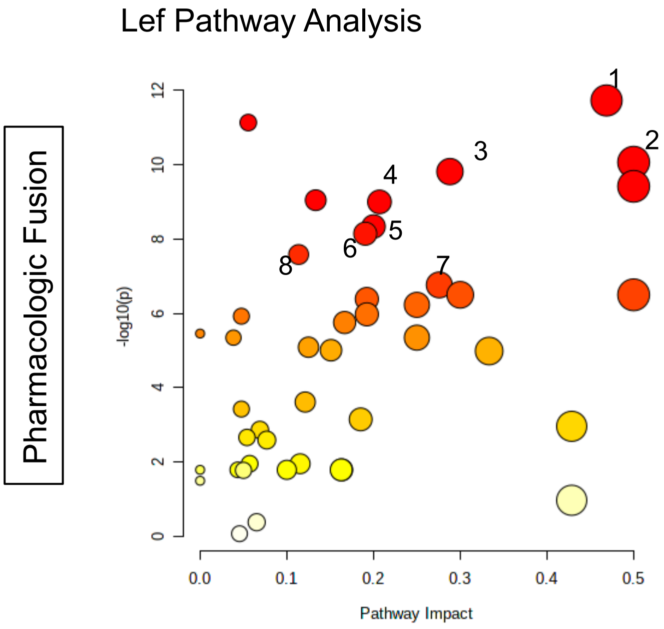


1. Alanine, Aspartate, and Glutamate Metabolism
2. Arginine Biosynthesis
3. Pyrimidine Metabolism
4. Aminoacyl t-RNA Metabolism
5. Arginine and Proline Metabolism
6. Nicotinate and Nicotinamide Metabolism
7. TCA Cycle
8. Purine Metabolism

**Figure S1.** Pathway analysis generated from discriminant metabolites identified by univariate Student's *t*-test.

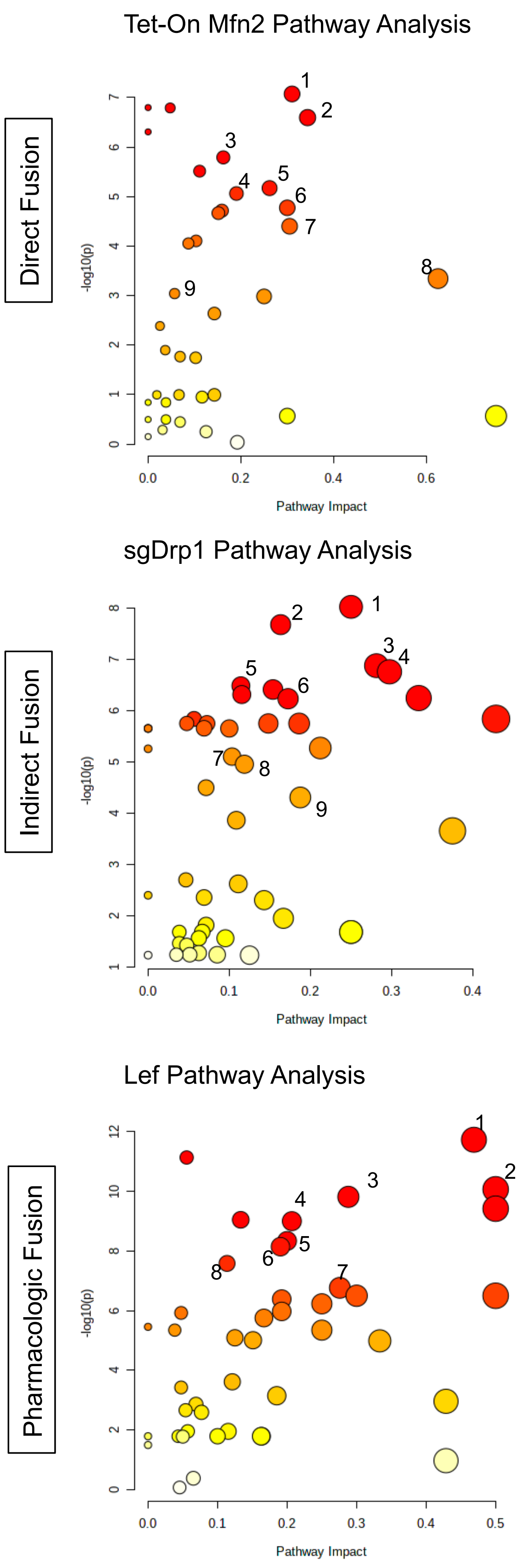


**Tet-On Mfn2 (Direct Fusion)**

1. Alanine, Aspartate, and Glutamate Metabolism
2. Arginine Biosynthesis
3. Pyrimidine Metabolism
4. Aminoacyl t-RNA Metabolism
5. Arginine and Proline Metabolism
6. Nicotinate and Nicotinamide Metabolism
7. TCA Cycle
8. Purine Metabolism

**sgDrp1 (Indirect Fusion)**

1. Arginine and Proline Metabolism
2. Fatty Acid Degradation
3. Alanine, Aspartate, and Glutamate Metabolism
4. Glutathione Metabolism
5. Glycolysis/Gluconeogenesis
6. TCA Cycle
7. Pentose Phosphate Pathway
8. Pyrimidine Metabolism
9. Arginine Biosynthesis

**Lef (Pharmacologic Fusion)**

1. Alanine, Aspartate, and Glutamate Metabolism
2. Arginine Biosynthesis
3. Pyrimidine Metabolism
4. Aminoacyl t-RNA Metabolism
5. Arginine and Proline Metabolism
6. Nicotinate and Nicotinamide Metabolism
7. TCA Cycle
8. Purine Metabolism

**Figure S2.** Pathway analysis generated from discriminant metabolites identified by SAM.

Direct Fusion

Tet-On Mfn2 Pathway Analysis

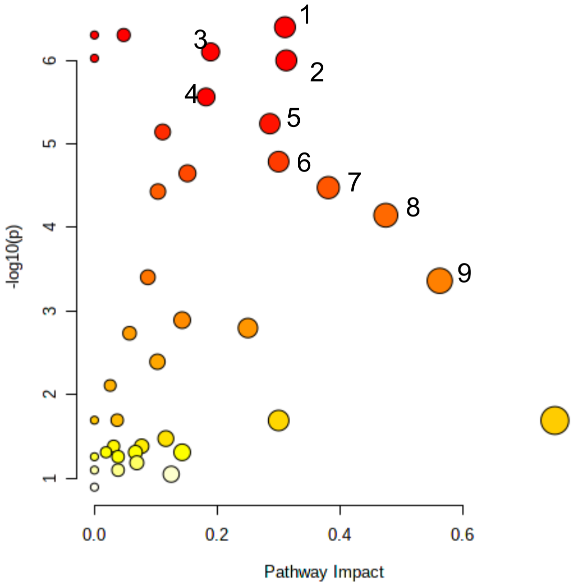


1. Aminoacyl-tRNA Biosynthesis
2. Alanine, Aspartate, and Glutamate Metabolism
3. Glutathione Metabolism
4. Purine Metabolism
5. Glycine, Serine, and Threonine Metabolism
6. Arginine and Proline Metabolism
7. Nicotinate and Nicotinamide Metabolism
8. Pyrimidine Metabolism
9. Arginine Biosynthesis

Indirect Fusion

sgDrp1 Pathway Analysis

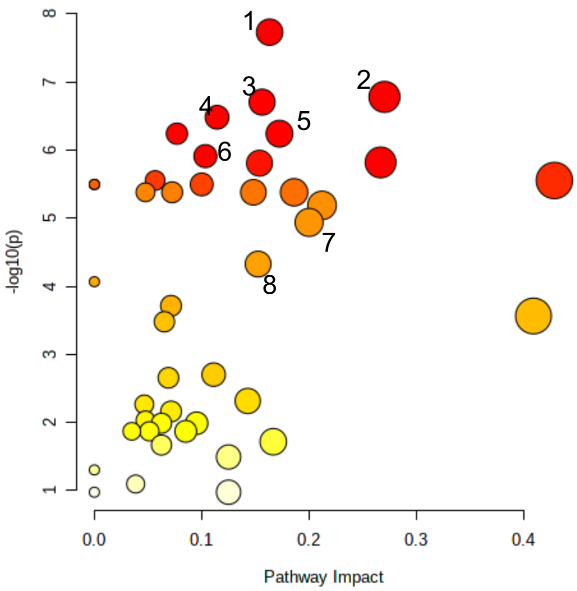


1. Fatty Acid Degradation
2. Glutathione Metabolism
3. Alanine, Aspartate, and Glutamate Metabolism
4. Glycolysis/Gluconeogenesis
5. TCA Cycle
6. Pentose Phosphate Pathway
7. Arginine and Proline Metabolism
8. Pyrimidine Metabolism

Pharmacologic Fusion

Lef Pathway Analysis

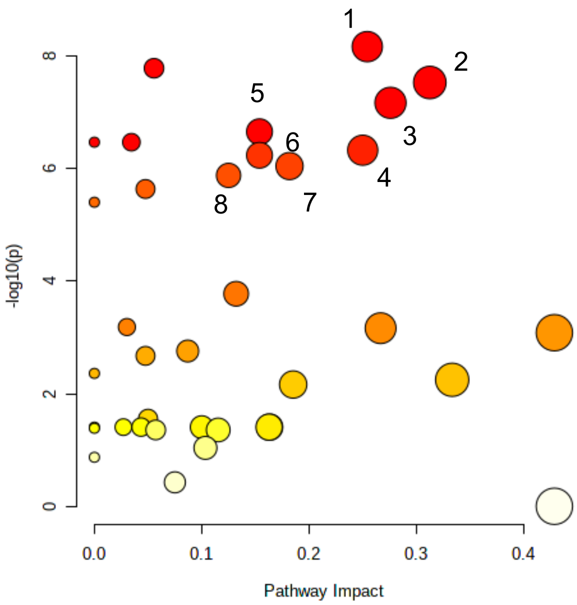


1. Pyrimidine Metabolism
2. Alanine, Aspartate, and Glutamate Metabolism
3. TCA Cycle
4. Valine, Leucine, and Isoleucine Biosynthesis
5. Glycoxylate and Dicarboxylate Metabolism
6. Glycerophospholipid Metabolism
7. Purine Metabolism
8. Arginine Biosynthesis

**Figure S3.** Pathway analysis generated from discriminant metabolites identified by PLS-DA VIP.

### Tet-On Mfn2 Pathway Analysis

Direct Fusion

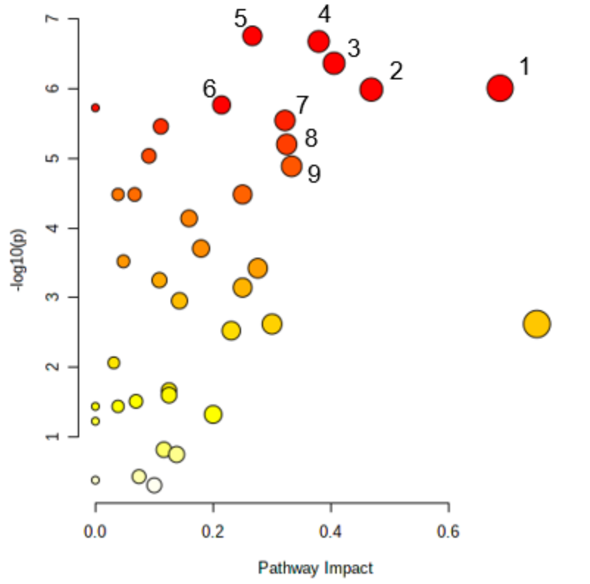


1. Arginine Biosynthesis
2. Alanine, Aspartate, and Glutamate Metabolism
3. Glutathione Metabolism
4. Aminoacyl-tRNA Biosynthesis
5. Histidine Metabolism
6. Glycine, Serine, and Threonine Metabolism
7. Pyrimidine Metabolism
8. Arginine and Proline Metabolism
9. Nicotinate and Nicotinamide Metabolism

### sgDrp1 Pathway Analysis

Indirect Fusion

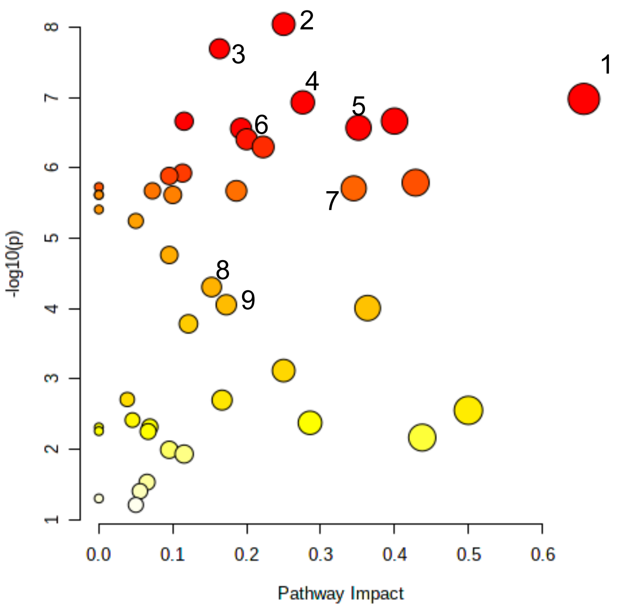


1. Alanine, Aspartate, and Glutamate Metabolism
2. Arginine and Proline Metabolism
3. Arginine Biosynthesis
4. TCA Cycle
5. Glutathione Metabolism
6. Glycolysis/Gluconeogenesis
7. Pentose Phosphate Pathway
8. Pyrimidine Metabolism
9. Aminoacyl-tRNA Biosynthesis

### Lef Pathway Analysis

Pharmacologic Fusion

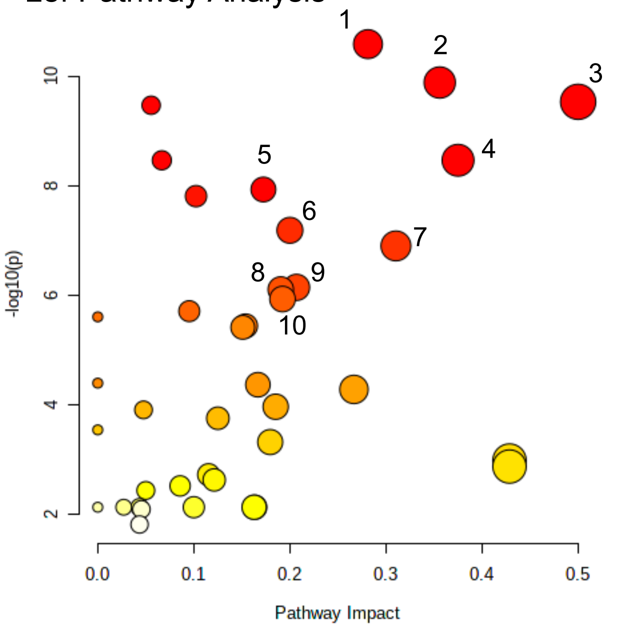


1. Alanine, Aspartate, and Glutamate Metabolism
2. Pyrimidine Metabolism
3. Valine, Leucine, and Isoleucine Biosynthesis
4. Arginine Biosynthesis
5. Aminoacyl t-RNA Metabolism
6. Arginine and Proline Metabolism
7. TCA Cycle
8. Pentose Phosphate Pathway
9. Nicotinate and Nicotinamide Metabolism
10. Glycerophospholipid Metabolism

**Figure S4.** Pathway analysis generated from discriminant metabolites identified by RF classification.

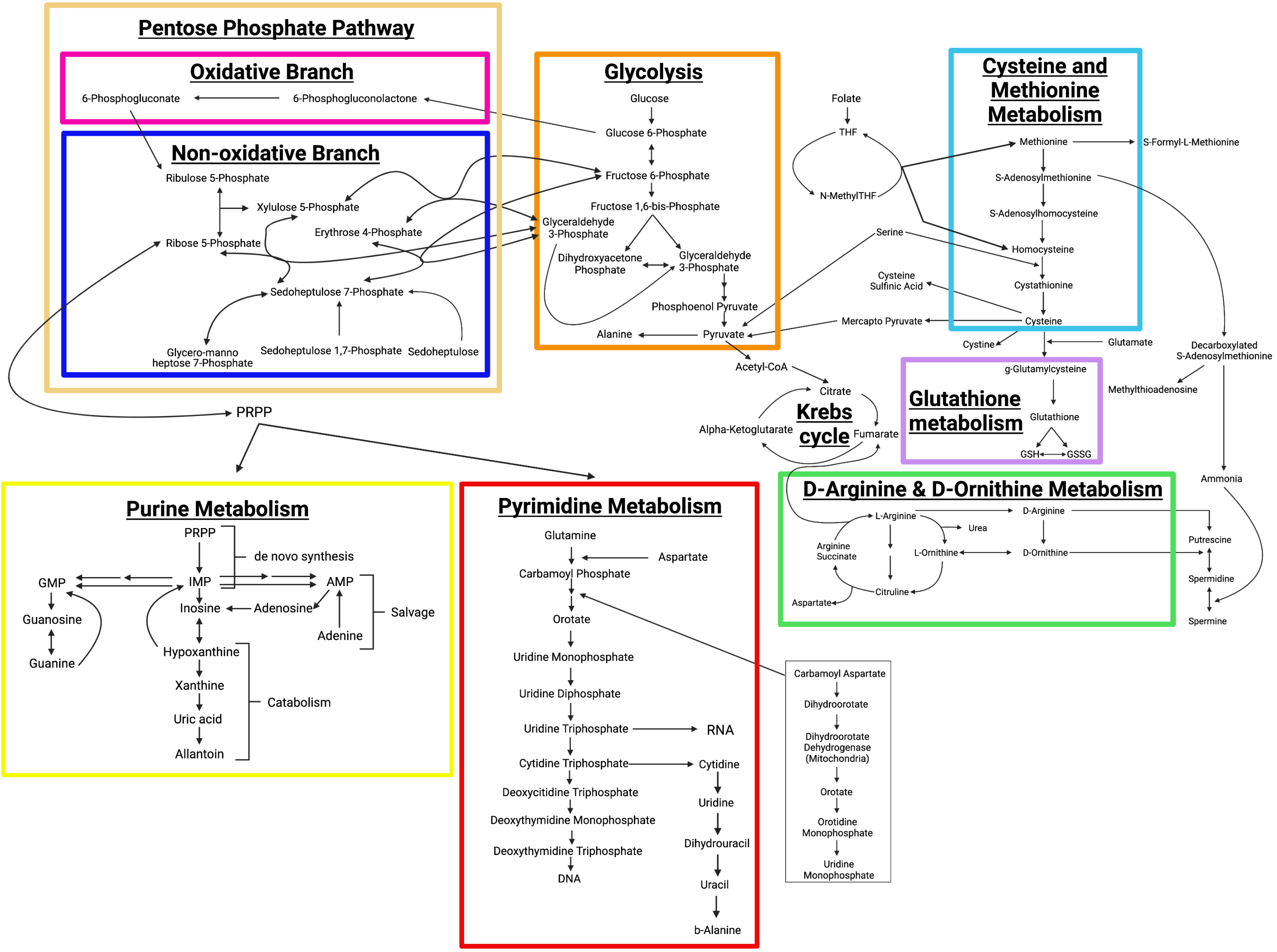


**Figure S5.** Significantly altered metabolic pathways in fusion induced KPC cells interconnected. Top 8 altered metabolic pathways identified from our overlapped discriminant metabolite set in fused KPC cells are mapped to show their interdependent relationships among each other.
